# Benchmarking kinetic models of *Escherichia coli* metabolism

**DOI:** 10.1101/2020.01.16.908921

**Authors:** Denis Shepelin, Daniel Machado, Lars K. Nielsen, Markus J. Herrgård

## Abstract

Predicting phenotype from genotype is the holy grail of quantitative systems biology. Kinetic models of metabolism are among the most mechanistically detailed tools for phenotype prediction. Kinetic models describe changes in metabolite concentrations as a function of enzyme concentration, reaction rates, and concentrations of metabolic effectors uniquely enabling integration of multiple omics data types in a unifying mechanistic framework. While development of such models for *Escherichia coli* has been going on for almost twenty years, multiple separate models have been established and systematic independent benchmarking studies have not been performed on the full set of models available. In this study we compared systematically all recently published kinetic models of the central carbon metabolism of *Escherichia coli*. We assess the ease of use of the models, their ability to include omics data as input, and the accuracy of prediction of central carbon metabolic flux phenotypes. We conclude that there is no clear winner among the models when considering the resulting tradeoffs in performance and applicability to various scenarios. This study can help to guide further development of kinetic models, and to demonstrate how to apply such models in real-world setting, ultimately enabling the design of efficient cell factories.

**Author summary:** Kinetic modeling is a promising method to predict cell metabolism. Such models provide mechanistic description of how concentrations of metabolites change in the cell as a function of time, cellular environment and the genotype of the cell. In the past years there have been several kinetic models published for various organisms. We want to assess how reliably models of *Escherichia coli* metabolism could predict cellular metabolic state upon genetic or environmental perturbations. We test selected models in the ways that represent common metabolic engineering practices including deletion and overexpression of genes. Our results suggest that all published models have tradeoffs and the model to use should be chosen depending on the specific application. We show in which cases users could expect the best performance from published models. Our benchmarking study should help users to make a better informed choice and also provides systematic training and testing dataset for model developers.

## Introduction

Reliable prediction of metabolic phenotypes in form of intracellular metabolite concentrations and metabolic fluxes from genotype would be a transformative technology for biotechnology and metabolic physiology. Instead of expensive and laborious screening experiments biologists could employ in silico tools to design cells that produce chemicals or proteins, or to design experiments that unravel additional complexity of metabolism. A plethora of metabolic phenotype prediction methods have been developed ranging from purely statistical machine learning approaches (1, 2) to mechanistic simulators of virtual cells (3). However, the complexity of living systems including the rich network of regulation that connects DNA sequence to the metabolic phenotype makes the phenotypic prediction task challenging.

Kinetic models are among the most mechanistically detailed methods for phenotype prediction, and they also have a long history of application for studying metabolism due to availability of in vitro kinetic parameter data for many enzymes in central carbon metabolism. These models are mathematical models describing changes in metabolite concentrations as a function of concentrations of enzymes catalyzing reactions, their kinetic parameters, and concentrations of substrates, products and other metabolic effectors (4). Usually such models are expressed in the form of ordinary differential equations (ODEs) and analyzed using methods that are suitable for solving and studying complex systems of ODEs. From the user perspective kinetic models can predict concentration profiles of metabolites and metabolic fluxes over time given some initial concentrations of metabolites and information on genetic perturbations such as deletion or overexpression of genes. Kinetic models are usually represented as a set of metabolic reactions, set of kinetic parameters inferred from *in vitro* or *in vivo* experimental data, and a set of initial concentrations of intracellular metabolites encoded together in the Systems Biology Markup Language (SBML) (5) format or as executable scripts for platform such as MATLAB.

Despite their long history and demonstrated ability to predict specific metabolic phenotypes used to for example parameterize models, there is a lack of systematic and unbiased evaluation studies of the predictive power of kinetic models (6, 7). The introduction of a comprehensive set of benchmarks covering different experimental datasets and use scenarios would allow to explore the limitations of existing kinetic models and continuously improve the models.

This study presents a systematic evaluation of the most representative kinetic models of *E. coli* central carbon metabolism. Models are compared in terms of their accuracy in predicting steady-state fluxes upon such perturbations as gene knockouts, changes in enzyme abundance, and changes of cultivation conditions. We provide Jupyter notebooks with simulations, code, and data, and encourage all model developers to test newly constructed models against the proposed set of benchmarks, and/or extend the benchmarks themselves with new experimental flux datasets.

## Methods

### Models

In this study we examine the most recent kinetic models of *E. coli* metabolism as well as an older widely-used kinetic model that many of the more recent models used as a starting point for development (Figure 1). In the following we describe basic statistics for the models (Table 1) and several other key model features such as distribution mechanism, use of standardized IDs for metabolites and reactions (Table 2).

**Fig 1.**
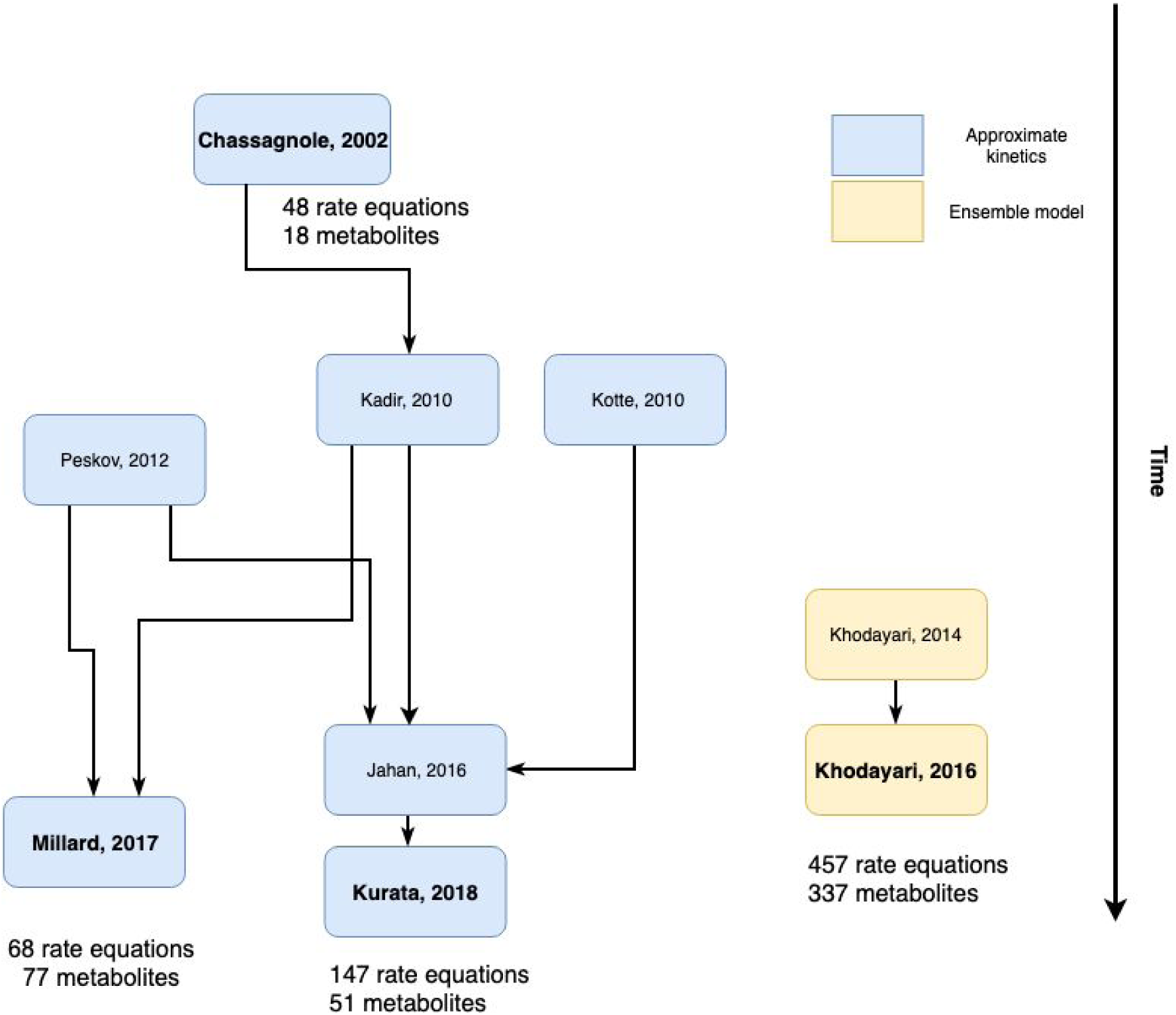
Lineage of *E. coli* dynamic models. Arrows represent at least partial usage of data/assumptions in the models. Models with the author names in bold are considered in the benchmark.

**Table 1.** Scope of benchmarked models. PTS - phosphotransferase system, EMP - Embden-Meyerhof-Parnas pathway, ED - Entner-Doudoroff pathway, PPP - Pentose Phosphate pathway, TCA - tricarboxylic acid cycle, Genome scale - all reactions from annotated genome.

| Model | Model size | Biochemical scope | Data used |
| --- | --- | --- | --- |
| Chassagnole02 | 48 rate equations<br>18 metabolites | PTS, EMP, PPP, biosynthesis of biomass precursors | (8) |
| Khodayari16 | 457 rate equations<br>337 metabolites | Genome scale | (9), (10,11) |
| Millard17 | 68 rate equations<br>77 metabolites | PTS, EMP, ED, PPP, TCA (with glyoxylate), acetate secretion | (12,13) |
| Kurata18 | 147 rate equations<br>51 metabolites | PTS, EMP, ED, PPP, TCA (with glyoxylate), acetate secretion, enzyme production | (14) |
| iML1515<br>(constraint-based model) | 2712 rates<br>1877 metabolites | Genome scale | NA |

**Table 2.** Distribution mechanism and technical details of models implementation.

| Model | Implementation | IDs | Change of conditions | API to change parameters |
| --- | --- | --- | --- | --- |
| Chassagnole02 | SBML | Custom | Glucose feed | Yes |
| Khodayari16 | MATLAB | BiGG | No | No |
| Millard17 | SBML | Custom | Glucose feed | Yes |
| Kurata18 | MATLAB | Custom | Yes | No |

Chassagnole02: This model is one of the first published kinetic models of *E. coli* central metabolism and it has been quite widely used (8). This model is represented as set of ODEs and distributed as an SBML file deposited on BioModels (15). In-house experimental metabolite level timecourse data were used to train the model.

Khodayari16: This model was specifically made to predict physiological parameters of cell and it is the largest model included in this comparison (16). The model is an update of previously published genome-scale kinetic model (17). This model was trained on the dataset compiled from published papers describing metabolism of E. coli in chemostat conditions. Reaction and metabolite identifiers are following BiGG (18) standard naming scheme, which enable the user to connect metabolite level measurements to standard identifiers that are used in experimental work.

Millard17: This model was originally developed as an answer to questions about metabolic regulation and not as a tool for prediction (19). It uses some formulations from (20) and employs published in vitro kinetic parameters deposited in databases like BRENDA. In-house data were used to train the model, and separate datasets (12, 13) were used for model evaluation, but not for fitting model parameters.

Kurata18: This model (21) was designed to be used as metabolic phenotype prediction tool and is an update from the model published by (22). The model was trained on batch fermentation data from (14). Users can change cultivation conditions via dilution rate (for continuous cultivation simulation) or set fermentation to batch regime.

This set of models is not exhaustive. The model from (23) was excluded due to limited coverage of metabolic pathways (for example there is no Pentose Phosphate Pathway). The model from (20) is notable because it’s one of the first highly detailed and models, but it is unavailable in public model repositories such as JWS Online or Biomodels.

We have also included the most recent constraint-based model of *E. coli* metabolism (24) to create a baseline for evaluating kinetic model prediction performance. Constraint-based models do not account for enzymatic rate equations or kinetic parameters, and can only be used for performing steady state flux balance analysis. However, due to their relative simplicity, ease of use and genome-wide coverage, these models can be considered as a first choice to predict strain phenotypes. For the constraint-based model we use a technique called linear minimization of metabolic adjustment (l*MOMA*) to simulate the effect of genetic perturbations. Other perturbation simulation methods for constraint-based models exist, but the purpose of the current study is not to compare these methods together and lMOMA has demonstrated good performance and is commonly used.

### Simulation setup

Chassagnole02: We used the Python Tellurium package (25) to simulate kinetic models that were in SBML format. The model was modified by 1) fixing stoichiometry in PTS system 2) extracellular volume, and 2) removing training-data specific equations for cofactors. To match simulation regimes of other models we simulated the model for 10.000 seconds. There is no way to change growth rate or dilution rate, so one simulation was used for every experimental condition. The Chassagnole02 model does not include the TCA cycle and ED pathway and the corresponding fluxes were fixed to be 0 in the results presented below (the results for model comparisons excluding the TCA cycle and ED pathway flux are found in the Supplementary Material).

Khodayari16: This model is implemented as a set of MATLAB scripts, and we have used the scripts from the original publications and their internal functions to simulate phenotypes. The model assumes that the amount of enzymes in the system can be modulated, so a knockout can be implemented by setting the amount of an enzyme to be zero. There is no documented way to change dilution rate or other cultivation condition. The output of the model is a time course of metabolite concentrations and fluxes.

Millard17: The model is distributed in SBML format and the simulations were performed in the Tellurium package. To match the simulation regime of other models we simulated the Millard17 model for 10.000 seconds without checking for the steady state. To match growth conditions one can manipulate the glucose feed parameter, but in the case of this model, we were not able to achieve a growth rate higher than 0.4 h-1 regardless of the glucose feed.

Kurata18: We used the MATLAB scripts from the original publication. The source code does not provide any functions to modify the model parameters, so we modified them directly in the source code. This model has explicit dilution factors or starting glucose concentration to support both chemostat and batch regimes.

iML1515: For simulation of the constraint-based model iML1515 with lMOMA we used cobrapy (26) and cameo packages (27). Chemostat culture is simulated as minimization of glucose uptake with constraint on growth to match the experimental dilution rate. For simulating knockout scenarios we additionally set the flux of the respective reaction to zero. For simulations with lMOMA, we first used the experimental data for the wild-type strain in defined reference condition. Next, we matched these data as close as possible to create a reference flux distribution for lMOMA, assuming that cells try to minimize flux changes upon knockout.

### Datasets

To compare accuracy of model predictions we gathered datasets describing physiology of *E. coli* grown on minimal medium with glucose with recorded metabolic flux profiles after various perturbations. Characteristics of datasets are provided in Table 3.

**Table 3.** Summary of metabolic flux dataset features (genotype, scope and experimental factors).

| Dataset | Sample genotypes | Fermentation | Number of fluxes |
| --- | --- | --- | --- |
| Ishii07 | WT, $\Delta galM$, $\Delta glk$, $\Delta pgm$, $\Delta pgi$, $\Delta pfkA$, $\Delta pfkB$, $\Delta fbp$, $\Delta fbaB$, $\Delta gapC$, $\Delta gpmA$, $\Delta gpmB$, $\Delta pykA$, $\Delta pykF$, $\Delta ppsA$, $\Delta zwf$, $\Delta pgl$, $\Delta gnd$, $\Delta rpe$, $\Delta rpiA$, $\Delta rpiB$, $\Delta tktA$, $\Delta tktB$, $\Delta talA$, $\Delta talB$ | Carbon-limited chemostat with glucose minimal media<br>Dilution = 0.2 h<sup>-1</sup> | 46 |
| Nicolas07 | WT, $\Delta zwf$, overexpression of *zwf* | Batch | 20 |
| Yao11 | WT | Chemostat<br>Dilution = 0.2, | 31 |
|  |  | 0.4, 0.6, 0.7<br>h-1 |  |
| Usui12 | WT, $\Delta pgi$, 4 levels of *pgi* expression,<br>4 levels of *eno* expression | Batch | 32 |
| Long19 | WT, $\Delta glk$, $\Delta pgm$, $\Delta pgi$, $\Delta pfkA$, $\Delta pfkB$,<br>$\Delta fbp$, $\Delta fbaB$, $\Delta tpiA$, $\Delta zwf$, $\Delta pgl$, $\Delta gnd$,<br>$\Delta rpe$, $\Delta sgcE$, $\Delta rpiB$, $\Delta tktA$, $\Delta tktB$,<br>$\Delta talA$, $\Delta talB$, $\Delta edd$, $\Delta eda$ | Batch | 79 |

For each scenario we converted identifiers of flux measurements to the BiGG ID nomenclature. Next, we calculated normalized fluxes by dividing each flux with the glucose update flux for each dataset. All datasets are available in the Supplementary Material in “tidy format” where each row represents one measurement (28).

### Scenarios

We propose to use three scenarios to compare the predictive performance of models:

1. Prediction of gene knockouts. In this scenario the models are used to predict the steady state fluxes of in silico mutants obtained by knocking out the reactions corresponding to a given gene. Data for this scenario is taken from ref. (11) for chemostat condition and from ref. (29) for batch condition. In this scenario we also perform constraint-based simulations using lMOMA to use as a baseline error estimate.
2. Prediction of flux profiles with respect to changes in enzyme abundance. In this scenario, the models are used to predict the steady-state fluxes of in silico mutants with varied abundance of single enzymes (glucose-6-phosphate dehydrogenase - *zwf*, glucose-6-phosphate isomerase - *pgi*, enolase - *eno*). Data for this scenario is taken from ref. (30) and ref. (31)
3. Prediction of flux profile with respect to changes in dilution rates. In this scenario, models are used to predict the steady-state fluxes of wild-type strains cultivated in chemostat with varied dilution rates. Data for this scenario is taken from ref. (32). In this scenario we also perform constraint-based simulations as baseline.

### Metrics

We employ two different metrics to assess the performance of each model. Each metric is calculated with fluxes normalized to the glucose uptake.

Firstly, we calculate “Normalized error” metric for each sample within scenario which is described in (33). This metric is calculated as:

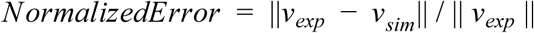

Where ||. || is an L2 norm, *v_exp_* is vector of fluxes defined in experimental conditions, *v_sim_* is vector of fluxes obtained per each sample in scenario.

Second, for each individual flux in sample we calculate “relative error” - absolute percentage error:

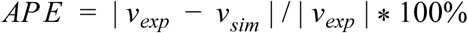

Where |.| is an absolute value, *v_exp_* is vector of fluxes defined in experimental conditions, *v_sim_* is vector of fluxes obtained per each sample in scenario. This metric is modified so 0 divided by 0 yields the 0% error. If either predicted or experimental data is zero, then the error is 100%.

## Results

We identified relevant kinetic models of *E. coli* metabolism through a literature search and selected models that covered major pathways in central-carbon metabolism and that were available in a usable format (either SBML or as MATLAB scripts). Most of the original publications describing the models did not systematically compare the new models to previously published models. This lack of assessment of model quality compared to previous models makes it hard to evaluate the choices made by modellers and to advance the field of kinetic modeling towards more predictive and comprehensive models.

Each model was trained on relatively small datasets, and these datasets were not common between the models. This fact allows us to perform out-of-sample tests of predictive qualities of the models using a larger dataset assembled from multiple independent publications. We selected experimental datasets that include 13C-MFA flux profiles (34) to test models. None of the experimental datasets had been used to train or fit the models except for the Khodayari16 model, which had been trained using the Ishii07 dataset. All the experimental data that is used in our benchmark is available as Table S1. Kinetic models can also be used to predict timecourses of metabolite concentrations, but we did not find enough experimental datasets containing time-course metabolomic profiles to build a sufficiently robust benchmark spanning different genotypes.

For each scenario (experiment) for each model we perform the following steps:

1) **Modify model parameters to match experimental conditions:** Select the parameters that are required to represent scenario (e.g. vmax = 0 for ZWF reaction to perform *zwf* knockout). Modify the selected model parameters. Set feed or dilution rate parameters to such value that model growth rate matches observed growth rate in the experiment as closely as possible.

2) **Simulate the model:** Simulate the model using the MATLAB files with custom scenario scripts for Khodayari16 and Kurata18 models, Tellurium for the Chassagnole02 and Millard17 models, and cobrapy for the constraint-based model.

3) **Post-process fluxes predicted by the model simulations:** Normalize fluxes to the glucose uptake. Select only such fluxes that are common between all models and the experimental dataset. Round normalized fluxes that are less than 0.1% of glucose uptake to be zero to avoid blowing up errors in small magnitude fluxes.

4) **Calculate prediction error metrics:** Metrics used were Absolute Percentage Error (APE) for individual fluxes in each sample, and normalized error for all the measured fluxes in a specific sample compared to the experimental dataset.

We simulate model performance in different scenarios described in the Methods Scenarios section.

### Simulation of gene knockout effects on fluxes

This scenario represents one of the most common experimental techniques in metabolic engineering. To perform gene knockout *in silico* in the model we force the parameter which corresponds to maximum velocity of reaction to be zero. Such qualitative modification may lead to the blocked reaction and failure to obtain phenotype as a result (Table 3A).

**Table 3A.** Phenotypes for which we were not able to simulate the model. Chemostat conditions.

| Model name | Genotype |
| --- | --- |
| Chassagnole02 | $\Delta fbp$, $\Delta pgl$, $\Delta pts$, $\Delta sucA$, $\Delta ppsA$, |
| Khodayari16 | $\Delta sdhCD$, $\Delta pts$ |
| Millard17 | $\Delta fbaA$, $\Delta fbaB$, $\Delta pfkA$, $\Delta pfkB$, $pts$, $\Delta sucA$, $\Delta tpi$ |
| Kurata18 | $\Delta gpmA$, $\Delta pgl$, $\Delta rpe$ |
| iML1515 | - |

**Table 3B.** Phenotypes for which we were not able to simulate the model. Batch condition.

| Model name | Genotype |
| --- | --- |
| Chassagnole02 | $\Delta fbp$, $\Delta pgl$, $\Delta pts$, $\Delta sucA$, $\Delta ppsA$, $eda$, $edd$ |
| Khodayari16 | $\Delta sdhCD$, $\Delta pts$ |
| Millard17 | $\Delta fbaA$, $\Delta fbaB$, $\Delta pfkA$, $\Delta pfkB$, $pts$, $\Delta sucA$, $\Delta tpi$ |
| Kurata18 | $\Delta gpmA$, $\Delta pgl$, $\Delta rpe$ |
| iML1515 | - |

Kinetic models Khodayari16 and Kurata17 are more accurate compared to constraint-based model iML1515 in chemostat setting (Fig 2A) and show comparable performance in batch setting (Fig 2B). Chassagnole04 model performs worse than iML1515 in chemostat condition and comparable performance in batch setting. We observe variability in the performance for individual models between various knockouts. Variability in performance does not seem to follow expectations that for reactions which are further from the start of the pathway the error in prediction would be higher. Besides general variability, some knockouts are outliers and these outliers are not common between models indicating some model implementation issues rather than complex nature of that particular knockout.

The Khodayari16 and Millard17 models show the best performance among kinetic models in the chemostat condition. We should note that Khodayari16 model was trained on the testing dataset which makes comparison to be biased for this model. The Chassagnole02 model has the worst performance across all kinetic models in all scenarios. This can be only partially attributed to lesser coverage of metabolic pathways such as TCA, Entner-Doudoroff pathway, acetate excretion (see Fig S1a and Fig S1b). Predictions depend on the model equations and availability of bypass pathways. Lack of bypass pathways often leads to model failing to capture the mutant phenotype at all resulting in simulation failure for example in case of Millard17 model (Table 3a and Table 3b). Authors of the Kurata18 model observed some numerical issues in simulating *pgl* knockout and such issues could also be related to implementation details of model equations.

Performance of models depend on cultivation regime used in experimental data. Both Khodayari16 and Millard17 models show good results in chemostat case and those models were trained on data from chemostat with dilution rate which is close to the dilution rate that was used in experimental dataset. The Kurata18 model was trained on batch culture data and it performs worse than the other models in the chemostat condition even if this model specifically allows for setting the cultivation condition to be chemostat. We see that the performance for Khodayari16 and Millard17 models drops when simulating batch physiology. None of the published models had been trained on both chemostat and batch data - doing so would most likely improve the prediction accuracy in both conditions.

Constraint based models generally perform quite poorly in case of pFBA, but methods such as lMOMA help to significantly improve their performance in cases where genetic perturbations are simulated. Accuracy of lMOMA simulation is just as good as kinetic models and in case of batch simulation even better (even if the predictions still have quite high errors). Ability to capture wild-type in experimental condition in the reference flux distribution seems to be exceptionally beneficial leading to improvements in prediction even without any inclusion of kinetic parameters in the model.

### Simulation of enzyme abundance variation on fluxes

This scenario highlights a unique feature of kinetic models, the ability to incorporate enzyme abundance data. Experimentally this aligns with the commonly used strategy to modulate gene expression in a specific pathway by replacing native promoters with constitutive or inducible heterologous promoters or by introducing additional copies of a gene on plasmids or by integrating them on the genome. Such scenarios cannot be easily represented using constraint-based models as the models only allow manipulating metabolic fluxes and not gene or protein expression. Instead the modeller would be required to use experimentally determined proteomic or proteomic data for the engineered strain together with one of the methods of transcriptomic or proteomic data integration (33). To simulate abundance of enzyme we multiply maximum velocity of reaction by a relative expression/abundance value obtained from corresponding study as result of qPCR assay. We assumed that changes in the transcript level directly translate into changes in the protein level.

Average accuracies for each model are not significantly different between each model and are significantly lower compared to performance of knockouts prediction in batch conditions. We could expect that the bigger the enzyme expression change is compared to WT the harder would be to predict flux profile, but we found no evidence for such pattern (Figure 3 and Figure S2).

**Fig 2a.**
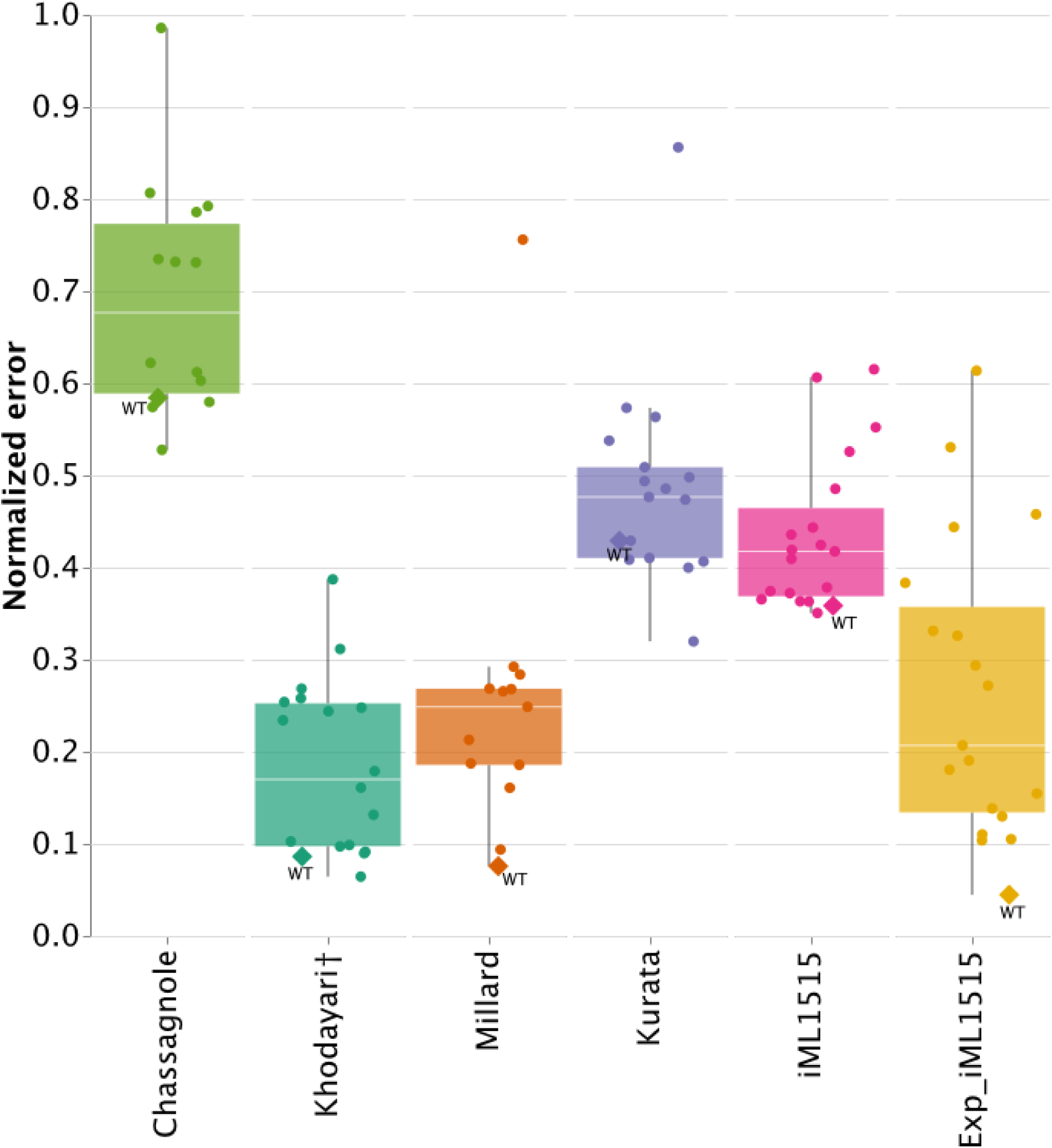
Error distribution for the gene knockout scenario in chemostat condition. Results are shown separately for the chemostat (A) and batch (B) simulations. Each dot represents a single comparison of simulation and experimental data (i.e. error across all measured fluxes for one particular strain/condition pair). Romb point shows prediction for wild-type sample. For boxplot the tick shows median, whisker spans from the smallest data to the largest data within the range [Q1 - k * IQR, Q3 + k * IQR] where Q1 and Q3 are the first and third quartiles while IQR is the interquartile range (Q3-Q1). The † symbol highlights that Khodayari16 model was trained on experimental dataset that is used in this chemostat knockouts scenario.

**Fig 2b.**
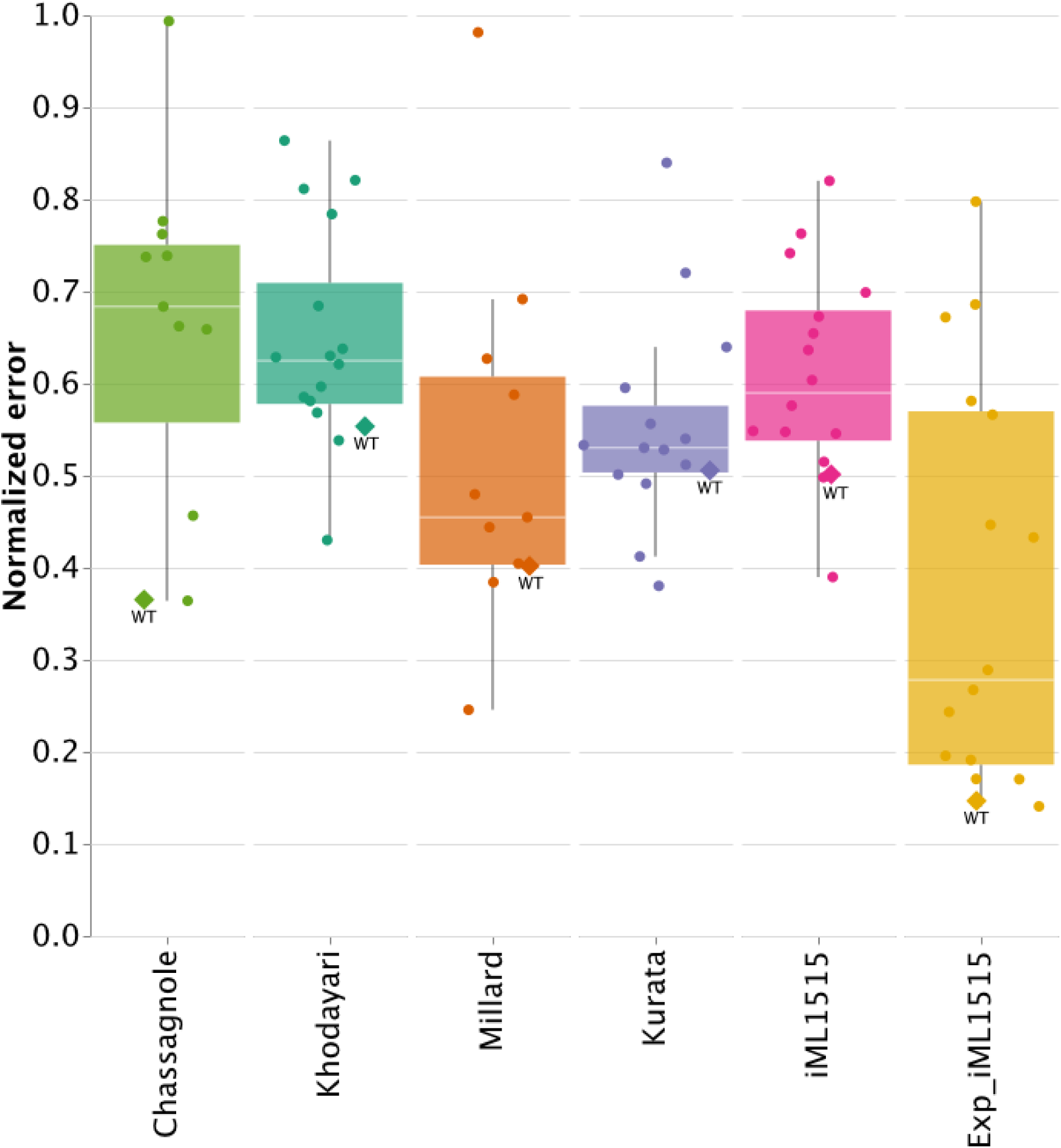
Error distribution for the gene knockout scenario in batch condition. Each dot represents a single comparison of simulation and experimental data. Box plots are drawn as explained in the caption for Figure 2a.

**Fig. 3.**
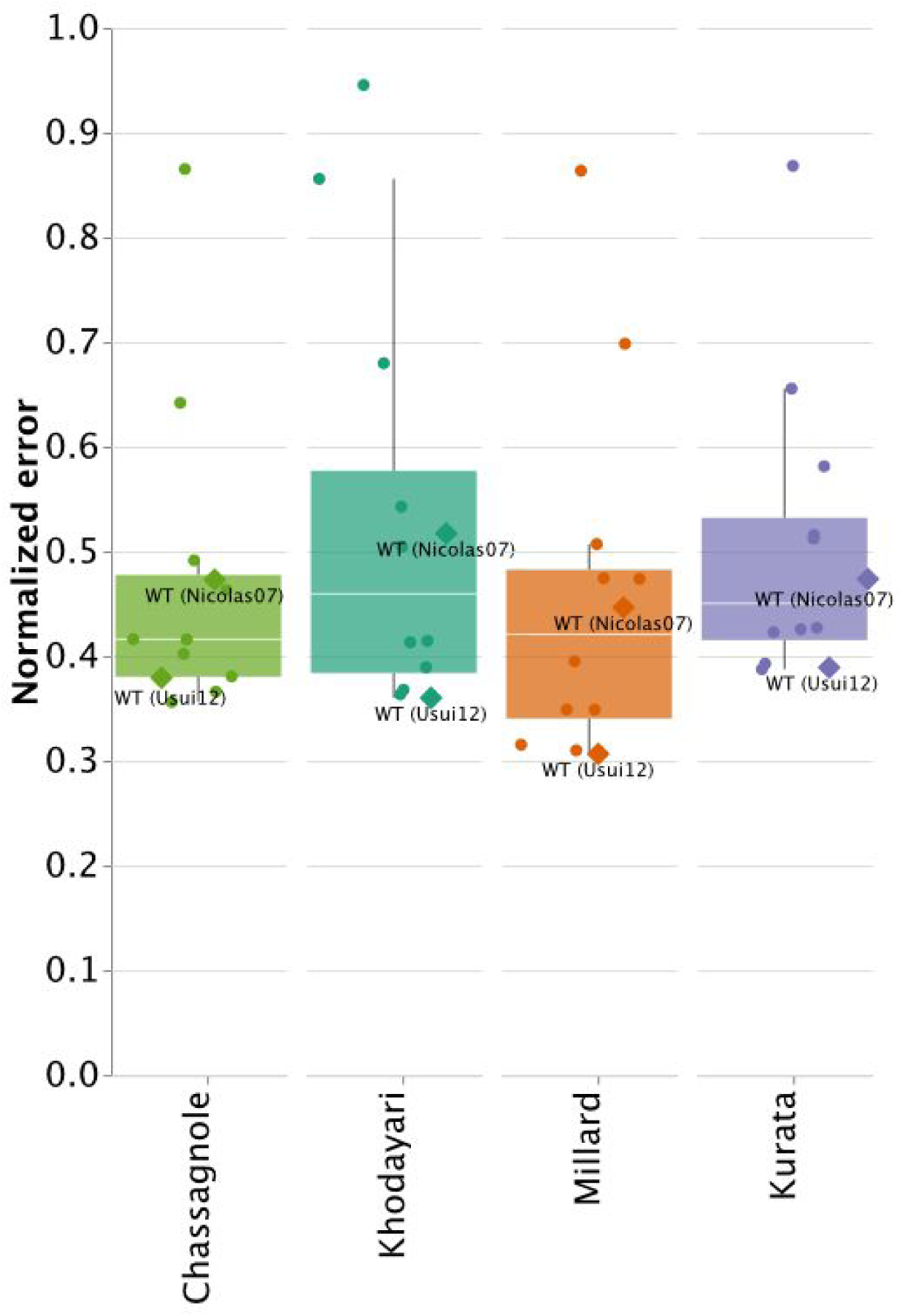
Errors distribution for the scenario “enzyme abundance”. Each dot represents a single comparison of simulation and experimental data. Box plots are drawn as explained in the caption for Figure 2a.

All of the models fail to predict the flux pattern for the 1.20-fold overexpression *pgi* strain (outlier data points in Figure 3). Despite a relatively modest increase in enzyme expression, the experimental data (31) indicates physiology of this strain differs from WT quite significantly. For example flux of reaction catalyzed by glucose-6-phosphate isomerase goes in gluconeogenesis direction (−5% of glucose uptake) in the engineered strain, while in WT it goes in glycolysis direction (60% of glucose uptake).

### Changes in dilution rate

The scenario with varied dilution rate tests the ability of kinetic models to react to the large-scale changes in the physiology of the cell. Compared to the previous scenarios there is no standard way to represent change in condition on parameter level for the models. Some models like Kurata18 allow direct manipulation by setting the dilution rate parameter, constraint-based models can be constrained to achieve growth rate equivalent to dilution rate, but most kinetic models have implicit assumption about allowed range of growth rates encoded in fitted parameter values. For example the Millard17 model is not capable of reaching a growth rate higher than 0.4 h-1 regardless of glucose feed flux.

Khodayari16 and Kurata18 model show expected behaviour - the closer experimental conditions are to the training dataset (0.2 h-1 chemostat in case of Khodayari16, batch with growth rate ≈ 0.7 h-1 in case of Kurata18) the higher the predictive accuracy is (Figure 4). The Chassagnole02 model that was trained on glucose pulses applied in a chemostat has the best predictive power for high dilution rates consistent with ability to handle conditions with high glucose uptake rate. Millard17 model is not able to reach growth rate more than 0.4 h-1 and we used simulation with growth rate of 0.4 h-1 to compare with experimental fluxes at 0.6 and 0.7 h-1. Surprisingly the error rate for those comparisons is similar to the dilution rates supported by the model natively.

**Fig. 4.**
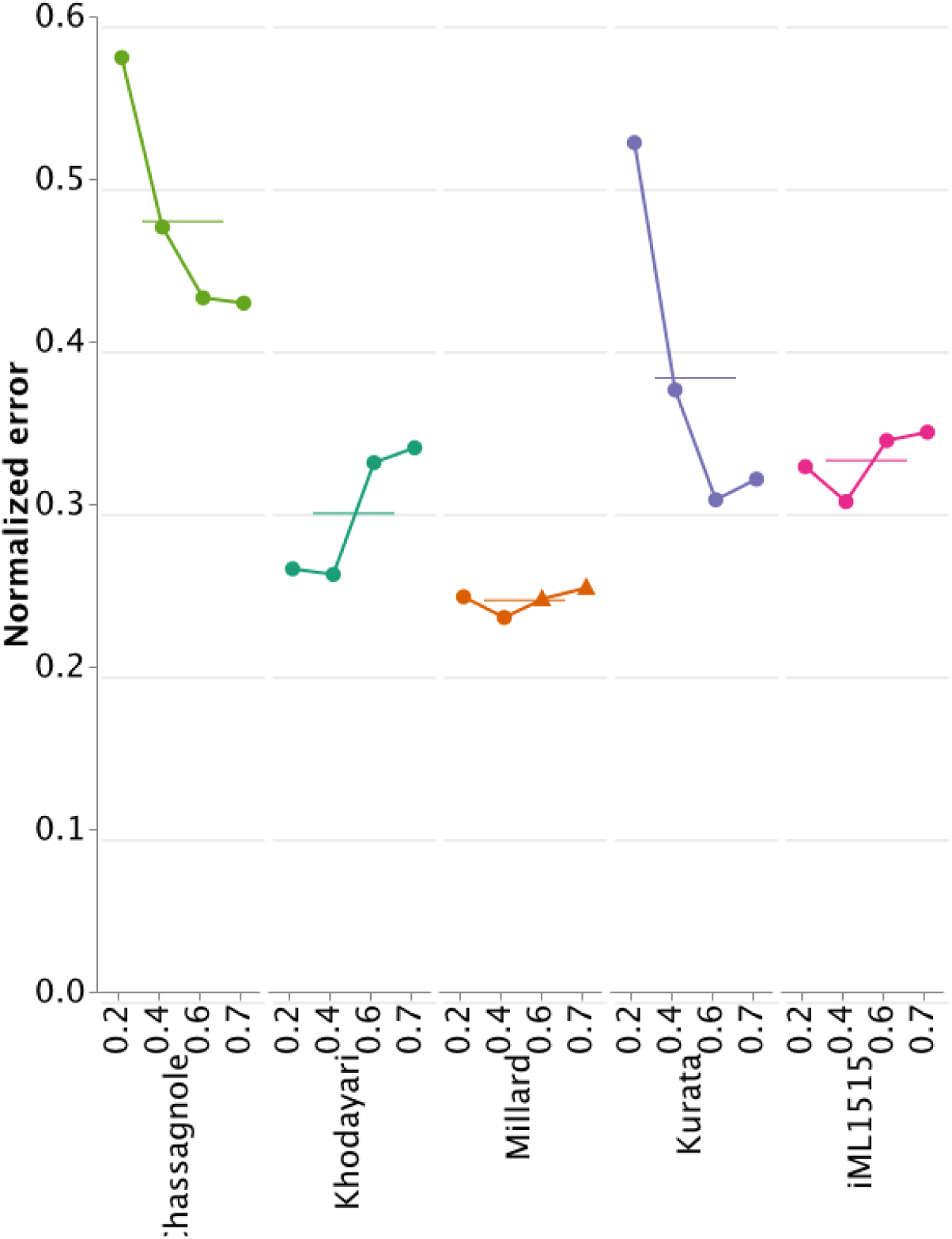
Errors distribution for the scenario “change in dilution rate”. Each dot represents a single comparison of simulation and experimental data. Ticks represent mean value. Triangles represent simulation of Millard17 model with a growth rate of 0.4 h-1 compared with experimental data at 0.6 and 0.7 h-1 dilution rates.

The constraint-based model shows a performance comparable to that of the kinetic models, with its accuracy degrading with higher dilution rates. This limitation was described earlier (33) and is attributed to the challenges of predicting activation of overflow metabolism without incorporating additional constraints (35).

## Discussion

We compared published kinetic models of E. coli metabolism and analyzed their predictive performance. Models were tested for their ability to predict steady-state flux profiles of knockout mutants, mutants with over/underexpression of enzyme and wild-type strain fluxes at different dilution rates. We found that none of the models were clearly better than others for all conditions. The older Chassagnole02 model has a smaller scope than the newer models and is worse compared to newer kinetic models and pFBA simulation.

There is now a growing body of evidence that systematic benchmarking studies stimulate research communities, help to set operating standards for evaluating computational models and methods, and lower the barriers for introducing new ideas in the field (36–38).

The development of kinetic models is presently an active field, in particular the automatic creation of kinetic models using predefined rules (39–42). However, most papers describing the development of new kinetic models do not include systematic benchmarking of the new model against existing ones. A previous publication presented a comparison of kinetic model performance (6), but since that time both new models and new experimental flux datasets have been published allowing significant extension of the benchmark. We propose a set of scenarios that are common in the metabolic engineering field - simulation of knockout strain and modulation of enzyme expression which could be used by authors of new models to perform unbiased assessment of their models. We have provided all the experimental data needed for benchmarking models with uniform set of identifiers to facilitate ease of use for future studies.

### Dynamics

Kinetic models are designed to capture dynamics. In practice, most models are trained and tested using steady-state flux and metabolite profiles. This situation is not greatly helped by shifting to batch cultures as these consist of a long period of near exponential growth (pseudo-steady state) followed by a brief transient and ultimate crash of the culture. Proper dynamic experiments would help resolve time constants for metabolic reactions and regulation (seconds), expression of enzymes (minutes), and cell growth and reactor dynamics (hours).

Unfortunately, dynamic metabolite level data is still not widely available. Fast dynamic data could potentially resolve kinetic parameters with much better precision, and information content in such studies would allow for the selection of the most appropriate functional form for kinetic rate laws (43). Availability of new datasets such as dynamic metabolite level data and increased biochemical knowledge can fuel the development of a new generation of models.

### Parametrization

Model parametrization is one area that impacts both model developers and users. With widespread use of omics technologies one can expect to see models adapting to use transcriptomic or proteomic (44) datasets for (re)calibration, but for most kinetic models users can not easily recalibrate parameters based on new data. An example of parameters that could be easily obtained experimentally are protein concentrations of metabolic enzymes. In the models included in our study protein concentrations are factored in the kinetic parameters such as maximal reaction rates. Such parametrization does not allow for change of enzyme abundances in an easy way even in situations where such data are available, because the original protein concentrations used to parameterize the models are not reported. Inclusion of parameters such as protein concentrations would enable the use of models in a wider set of conditions if users would know the protein abundance for the strain in given condition (45). Explicit absolute enzyme abundance is a good candidate for inclusion in the model parameter space as such parameters should allow adaptation of models to new conditions and non-standard genetic backgrounds.

### Getting the baseline right

The highly regulated nature of metabolism poses challenges in predicting which perturbations would create changes in the flux profile and which perturbations could be smoothed by homeostasis. In such a case the simplest null-model is an assumption that fluxes simply do not change at all between conditions or that change would be minimal. In our benchmark, we see that models make imperfect predictions for the basic metabolic state.

Success of lMOMA simulation highlights that even very simple models benefit a lot from factoring in baseline strain data. The ability to predict the behaviour of the wild-type strain in a given condition is an important property.

We can expect that inclusion of a broader spectrum of conditions for wild-type *E. coli* in training data would improve accuracy. Prediction errors could be decomposed as a sum of prediction error of wild-type in given condition simulation plus contribution of perturbation of the WT. Additional information about WT physiology in condition would improve that part of error.

### Uncertainty

All analyzed models work in the deterministic regime - there is no uncertainty in predictions. In the case of complex non-linear systems, it is hard for users to evaluate how sensitive the system is to small perturbations such as errors in experimental data. Most experimental research is now reported with at least some notion of uncertainty (46). Ideally models should propagate uncertainty from training data and parameters to the actual predictions (47, 48). Some kinetic frameworks try to perform uncertainty quantification in form of Monte Carlo sampling (49) or using a full Bayesian framework (50), but there is no published E. coli kinetic model using these frameworks that we could test in our benchmark. We hope that uncertainty quantification would be integrated into the next generation of models enabling exploration of the whole set of predictions rather than only the point estimates.

It should be noted that the reported flux values are inferred from mass-spectrometry measurements according to some measurement flux model. The closer the measurement flux model is to the kinetic model the better would be the predictions due to purely stoichiometric reasons. This may be a source of some bias in our assessment, but re-estimating fluxes using different measurement flux models is well beyond the scope of the current study.

### Complexity vs Accuracy

Models in the benchmark have different tradeoffs in scope versus biochemical details. Kurata18 and Millard17 model use detailed and complex kinetic rate laws (Monod-Wyman-Changeux) that allow a full and realistic description of allostery. Due to the complexity of the rate laws, the scope of the models is limited to a smaller part of the whole metabolism (Table 2), but on the other hand known allosteric interactions can be included in the model. On the other hand the Khodayari16 model uses universal Michaelis-Menten-like kinetic laws for every reaction with their ensemble approach. The use of such universal kinetic laws allows the building of a model with broader scope, but on the other hand there is less flexibility to represent known complexity in enzyme mechanisms. Constraint-based models of course represent the extreme version of models with a broad scope, but no actual kinetic rate laws.

### Standards

Usage of standard identifiers for metabolites and reaction that could be mapped to databases such as KEGG or MetaCyc should be enforced by the community. Arbitrary naming systems hinder comparison of models and integration with published data. For various reasons, most of the models included in this benchmark were not distributed in standard formats such as SBML. In some cases it may be justified due to incompleteness of format (ensemble model representation for example), but usage of SBML benefits both model builders and users. A rich set of tools for simulation, analysis and manipulation of SBML models exists and authors can rely on these tools to create a better user experience. Besides such technical benefits, SBML models are more easily discoverable due to existence of catalogs like BioModels or JWS that provide structured search and even web interface to simulation results. We hope that in the future more researchers will opt for representing their models in SBML format and hence ensure that their work finds its way to practical applications in metabolic engineering and other fields.

## Conclusion

Our benchmark highlighted how complex the task of accurate prediction of metabolic physiology in response to genetic and environmental perturbations is even for a simple and well characterized bacterium like E. coli. Mechanistic models for such complex task are also not expected to be simple. Model developers need to combine deep biochemical knowledge to describe the system, computer science and mathematical skills to select the right implementation formalism, and also a solid understanding of caveats in experimental data used to parameterize models (e.g. *in vitro* kinetics and metabolomics). We analyzed how decisions made by model developers impact accuracy of prediction across our whole benchmark dataset. We found that the mathematical formalism and kinetic complexity is less important for the ability to make accurate predictions than inclusion of wider set of conditions for model training. Models that were trained on very limited number of datasets could in general not be used reliably to make predictions in conditions that were not included in the training dataset.

We should note that tools and methodologies for modelling do improve and mature. We are sure that the recent rapid developments in machine learning and in particular methods that fuse machine learning and mechanistic model will fuel progress and application of kinetic models in bioscience.

Despite progress in the modeling field and the availability of more data for training models some challenges still remain. Big perturbations in physiology such as change in the mode of fermentation is hard to model. Despite advances in standardizing the formats for modelling there is still room for improvement in the formats and for increased adoption of existing formats. Prediction uncertainty quantification is being tackled right now, but is yet to be widely used. Discoverability of experimental data to be trained on could be improved, as most of the datasets were published before the FAIR initiatives (51). To address this issue we combined published experimental data and harmonized it to be easier used for future model training or evaluation. All data and the notebooks with simulations and analysis of the models are available on Github https://github.com/biosustain/EcoliKineticBenchmark (https://doi.org/10.5281/zenodo.3610129).

## Acknowledgments

We would like to thank Lyubov Golubeva and Mikhail Shupletsov for valuable input on performance and user experience assessment of Khodayari model.

## Supplementary

**Fig S1a.**
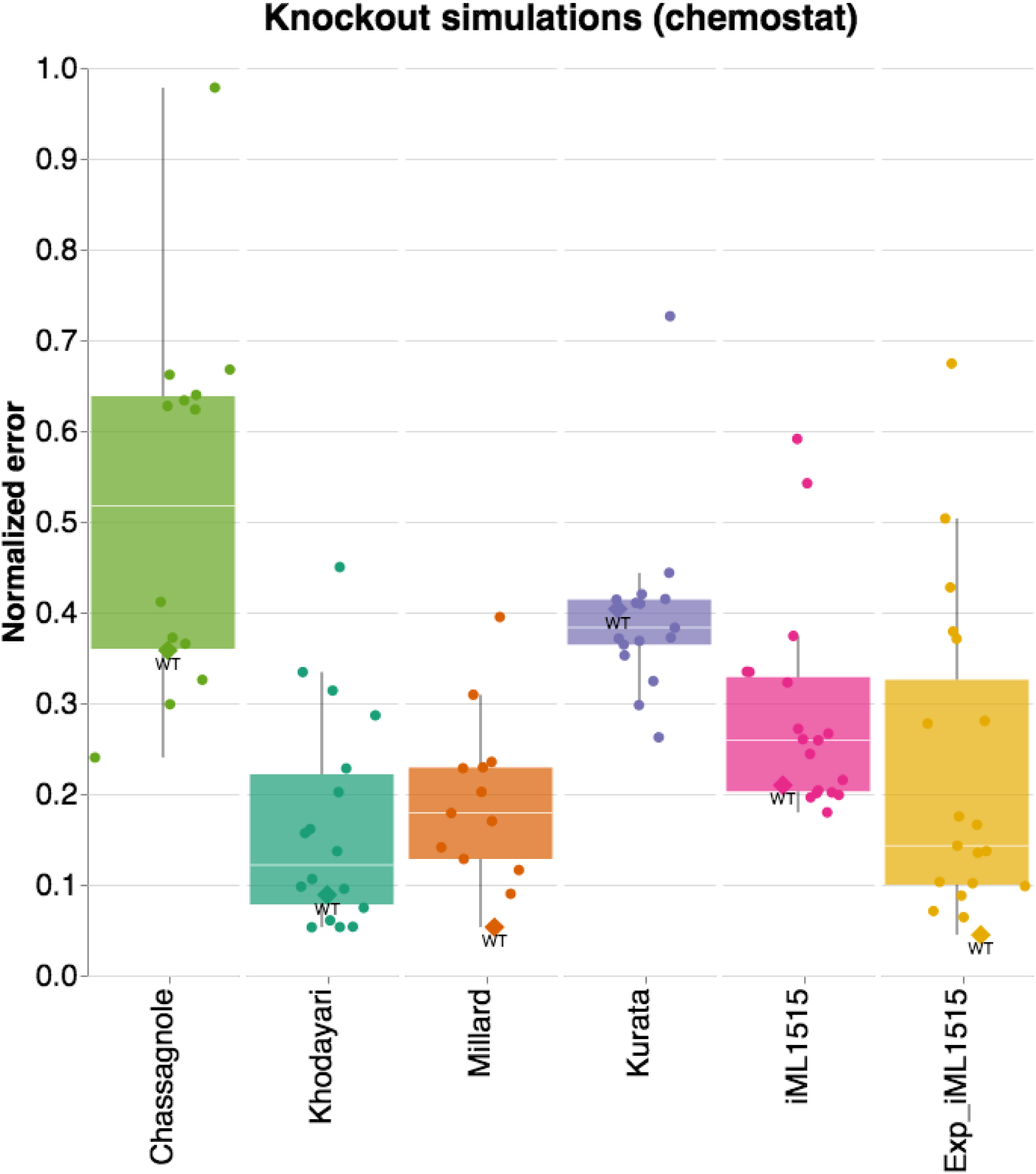
Errors in knockout prediction (chemostat) (only fluxes presented in Chassagnole02)

**Fig S1b.**
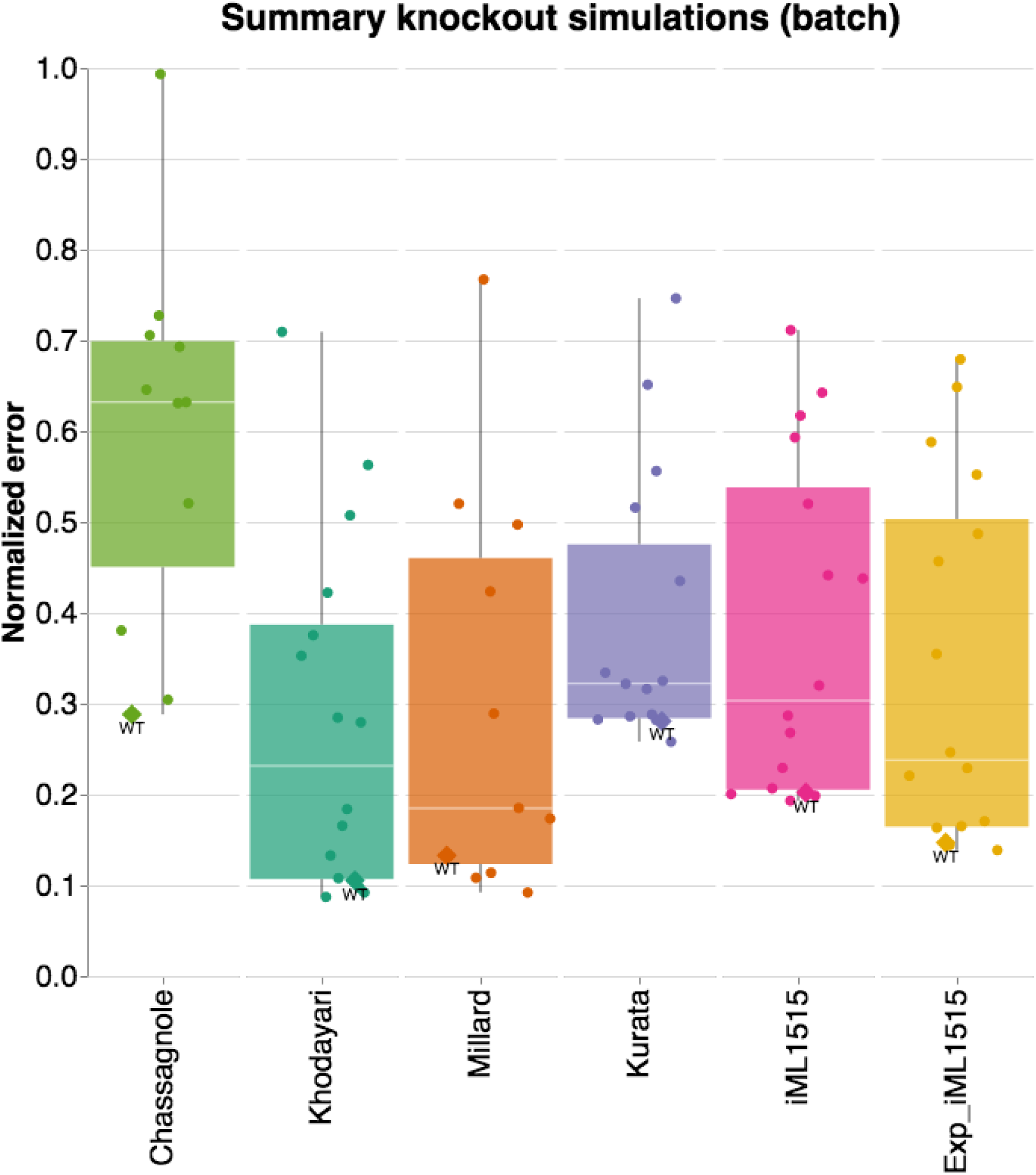
Errors in knockout prediction (batch) (only fluxes presented in Chassagnole02)

**Fig S2.**
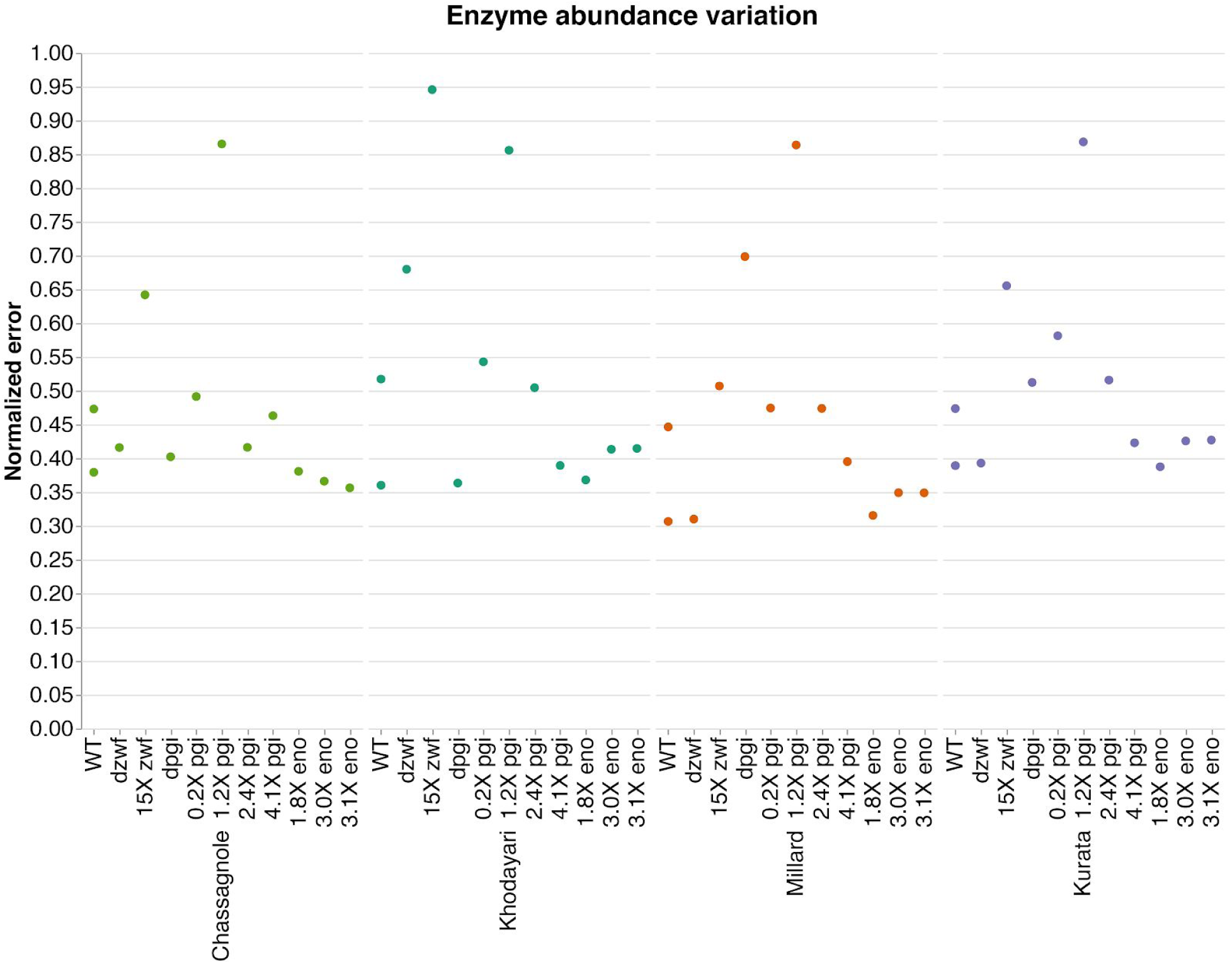
Normalized error enzyme abundance with individual samples.

## References

1. Costello Z, Martin HG. A machine learning approach to predict metabolic pathway dynamics from time-series multiomics data. npj Syst Biol Appl. 2018 May 29;4:19.

2. Zampieri G, Vijayakumar S, Yaneske E, Angione C. Machine and deep learning meet genome-scale metabolic modeling. PLoS Comput Biol. 2019 Jul 11;15(7):e1007084.

3. Karr JR, Sanghvi JC, Macklin DN, Gutschow MV, Jacobs JM, Bolival B, et al. A whole-cell computational model predicts phenotype from genotype. Cell. 2012 Jul 20;150(2):389–401.

4. Saa PA, Nielsen LK. Formulation, construction and analysis of kinetic models of metabolism: A review of modelling frameworks. Biotechnol Adv. 2017 Dec;35(8):981–1003.

5. Hucka M, Finney A, Sauro HM, Bolouri H, Doyle JC, Kitano H, et al. The systems biology markup language (SBML): a medium for representation and exchange of biochemical network models. Bioinformatics. 2003 Mar 1;19(4):524–31.

6. Lima AP, Baixinho V, Machado D, Rocha I. A Comparative Analysis of Dynamic Models of the Central Carbon Metabolism of Escherichia coli. IFAC-PapersOnLine. 2016;49(26):270–6.

7. Costa RS, Vinga S. Assessing Escherichia coli metabolism models and simulation approaches in phenotype predictions: Validation against experimental data. Biotechnol Prog. 2018 Oct 7;34(6):1344–54.

8. Chassagnole C, Noisommit-Rizzi N, Schmid JW, Mauch K, Reuss M. Dynamic modeling of the central carbon metabolism of Escherichia coli. Biotechnol Bioeng. 2002 Jul 5;79(1):53–73.

9. Zhao J, Baba T, Mori H, Shimizu K. Global metabolic response of Escherichia coli to gnd or zwf gene-knockout, based on 13C-labeling experiments and the measurement of enzyme activities. Appl Microbiol Biotechnol. 2004 Mar;64(1):91–8.

10. Kabir MM, Ho PY, Shimizu K. Effect of ldhA gene deletion on the metabolism of Escherichia coli based on gene expression, enzyme activities, intracellular metabolite concentrations, and metabolic flux distribution. Biochem Eng J. 2005 Nov;26(1):1–11.

11. Ishii N, Nakahigashi K, Baba T, Robert M, Soga T, Kanai A, et al. Multiple high-throughput analyses monitor the response of E. coli to perturbations. Science. 2007 Apr 27;316(5824):593–7.

12. Nanchen A, Schicker A, Sauer U. Nonlinear dependency of intracellular fluxes on growth rate in miniaturized continuous cultures of Escherichia coli. Appl Environ Microbiol. 2006 Feb;72(2):1164–72.

13. Taymaz-Nikerel H, van Gulik WM, Heijnen JJ. Escherichia coli responds with a rapid and large change in growth rate upon a shift from glucose-limited to glucose-excess conditions. Metab Eng. 2011 May;13(3):307–18.

14. Toya Y, Ishii N, Nakahigashi K, Hirasawa T, Soga T, Tomita M, et al. 13C-metabolic flux analysis for batch culture of Escherichia coli and its Pyk and Pgi gene knockout mutants based on mass isotopomer distribution of intracellular metabolites. Biotechnol Prog. 2010 Aug;26(4):975–92.

15. Glont M, Nguyen TVN, Graesslin M, Hälke R, Ali R, Schramm J, et al. BioModels: expanding horizons to include more modelling approaches and formats. Nucleic Acids Res. 2018 Jan 4;46(D1):D1248–53.

16. Khodayari A, Maranas CD. A genome-scale Escherichia coli kinetic metabolic model k-ecoli457 satisfying flux data for multiple mutant strains. Nat Commun. 2016 Dec 20;7:13806.

17. Khodayari A, Zomorrodi AR, Liao JC, Maranas CD. A kinetic model of Escherichia coli core metabolism satisfying multiple sets of mutant flux data. Metab Eng. 2014 Sep;25:50–62.

18. King ZA, Lu J, Dräger A, Miller P, Federowicz S, Lerman JA, et al. BiGG Models: A platform for integrating, standardizing and sharing genome-scale models. Nucleic Acids Res. 2016 Jan 4;44(D1):D515–22.

19. Millard P, Smallbone K, Mendes P. Metabolic regulation is sufficient for global and robust coordination of glucose uptake, catabolism, energy production and growth in Escherichia coli. PLoS Comput Biol. 2017 Feb 10;13(2):e1005396.

20. Peskov K, Mogilevskaya E, Demin O. Kinetic modelling of central carbon metabolism in Escherichia coli. FEBS J. 2012 Sep;279(18):3374–85.

21. Kurata H, Sugimoto Y. Improved kinetic model of Escherichia coli central carbon metabolism in batch and continuous cultures. J Biosci Bioeng. 2018 Feb;125(2):251–7.

22. Jahan N, Maeda K, Matsuoka Y, Sugimoto Y, Kurata H. Development of an accurate kinetic model for the central carbon metabolism of Escherichia coli. Microb Cell Fact. 2016 Jun 21;15(1):112.

23. Kotte O, Zaugg JB, Heinemann M. Bacterial adaptation through distributed sensing of metabolic fluxes. Mol Syst Biol. 2010 Mar 9;6:355.

24. Monk JM, Lloyd CJ, Brunk E, Mih N, Sastry A, King Z, et al. iML1515, a knowledgebase that computes Escherichia coli traits. Nat Biotechnol. 2017 Oct 11;35(10):904–8.

25. Choi K, Medley JK, König M, Stocking K, Smith L, Gu S, et al. Tellurium: An extensible python-based modeling environment for systems and synthetic biology. BioSystems. 2018 Sep;171:74–9.

26. Ebrahim A, Lerman JA, Palsson BO, Hyduke DR. COBRApy: COnstraints-Based Reconstruction and Analysis for Python. BMC Syst Biol. 2013 Aug 8;7:74.

27. Cardoso JGR, Jensen K, Lieven C, Lærke Hansen AS, Galkina S, Beber M, et al. Cameo: A python library for computer aided metabolic engineering and optimization of cell factories. ACS Synth Biol. 2018 Apr 20;7(4):1163–6.

28. Wickham H. Tidy Data. J Stat Softw. 2014;59(10).

29. Long CP, Antoniewicz MR. Metabolic flux responses to deletion of 20 core enzymes reveal flexibility and limits of E. coli metabolism. Metab Eng. 2019 Aug 4;

30. Nicolas C, Kiefer P, Letisse F, Krömer J, Massou S, Soucaille P, et al. Response of the central metabolism of Escherichia coli to modified expression of the gene encoding the glucose-6-phosphate dehydrogenase. FEBS Lett. 2007 Aug 7;581(20):3771–6.

31. Usui Y, Hirasawa T, Furusawa C, Shirai T, Yamamoto N, Mori H, et al. Investigating the effects of perturbations to pgi and eno gene expression on central carbon metabolism in Escherichia coli using (13)C metabolic flux analysis. Microb Cell Fact. 2012 Jun 21;11:87.

32. Yao R, Hirose Y, Sarkar D, Nakahigashi K, Ye Q, Shimizu K. Catabolic regulation analysis of Escherichia coli and its crp, mlc, mgsA, pgi and ptsG mutants. Microb Cell Fact. 2011 Aug 11;10:67.

33. Machado D, Herrgård M. Systematic evaluation of methods for integration of transcriptomic data into constraint-based models of metabolism. PLoS Comput Biol. 2014 Apr 24;10(4):e1003580.

34. Zamboni N, Fendt S-M, Rühl M, Sauer U. (13)C-based metabolic flux analysis. Nat Protoc. 2009 May 21;4(6):878–92.

35. de Groot DH, Lischke J, Muolo R, Planqué R, Bruggeman FJ, Teusink B. The common message of constraint-based optimization approaches: overflow metabolism is caused by two growth-limiting constraints. BioRxiv. 2019 Jun 21;

36. Boutros PC, Margolin AA, Stuart JM, Califano A, Stolovitzky G. Toward better benchmarking: challenge-based methods assessment in cancer genomics. Genome Biol. 2014 Sep 17;15(9):462.

37. Mangul S, Martin LS, Hill BL, Lam AK-M, Distler MG, Zelikovsky A, et al. Systematic benchmarking of omics computational tools. Nat Commun. 2019 Mar 27;10(1):1393.

38. Weber LM, Saelens W, Cannoodt R, Soneson C, Hapfelmeier A, Gardner PP, et al. Essential guidelines for computational method benchmarking. Genome Biol. 2019 Jun 20;20(1):125.

39. Gopalakrishnan S, Dash S, Maranas C. K-FIT: An accelerated kinetic parameterization algorithm using steady-state fluxomic data. BioRxiv. 2019 Apr 18;

40. St. John P, Strutz J, Broadbelt LJ, Tyo KEJ, Bomble YJ. Bayesian inference of metabolic kinetics from genome-scale multiomics data. BioRxiv. 2018 Oct 22;

41. Smith RW, van Rosmalen RP, Martins Dos Santos VAP, Fleck C. DMPy: a Python package for automated mathematical model construction of large-scale metabolic systems. BMC Syst Biol. 2018 Jun 19;12(1):72.

42. Miskovic L, Beal J, Moret M, Hatzimanikatis V. Model classification for uncertainty reduction in biochemical kinetic models. BioRxiv. 2018 Sep 27;

43. Vasilakou E, Machado D, Theorell A, Rocha I, Nöh K, Oldiges M, et al. Current state and challenges for dynamic metabolic modeling. Curr Opin Microbiol. 2016 Oct;33:97–104.

44. Schmidt A, Kochanowski K, Vedelaar S, Ahrné E, Volkmer B, Callipo L, et al. The quantitative and condition-dependent Escherichia coli proteome. Nat Biotechnol. 2016 Jan;34(1):104–10.

45. Sánchez BJ, Zhang C, Nilsson A, Lahtvee P-J, Kerkhoven EJ, Nielsen J. Improving the phenotype predictions of a yeast genome-scale metabolic model by incorporating enzymatic constraints. Mol Syst Biol. 2017 Aug 3;13(8):935.

46. Theorell A, Leweke S, Wiechert W, Nöh K. To be certain about the uncertainty: Bayesian statistics for 13 C metabolic flux analysis. Biotechnol Bioeng. 2017 Aug 23;114(11):2668–84.

47. Eriksson O, Jauhiainen A, Maad Sasane S, Kramer A, Nair AG, Sartorius C, et al. Uncertainty quantification, propagation and characterization by Bayesian analysis combined with global sensitivity analysis applied to dynamical intracellular pathway models. Bioinformatics. 2018 Jul 13;35(2):284–92.

48. Dass SC. Bayesian Solution Uncertainty Quantification for Differential Equations. Bayesian Anal. 2016 Dec;11(4):1275–7.

49. Miskovic L, Béal J, Moret M, Hatzimanikatis V. Uncertainty reduction in biochemical kinetic models: Enforcing desired model properties. PLoS Comput Biol. 2019 Aug 20;15(8):e1007242.

50. Saa PA, Nielsen LK. Construction of feasible and accurate kinetic models of metabolism: A Bayesian approach. Sci Rep. 2016 Jul 15;6:29635.

51. Wilkinson MD, Dumontier M, Aalbersberg IJJ, Appleton G, Axton M, Baak A, et al. The FAIR Guiding Principles for scientific data management and stewardship. Sci Data. 2016 Mar 15;3:160018.

